# GATA transcription factors act as gatekeepers between the Wnt/β-catenin-mediated proliferation and differentiation in the intestinal epithelium

**DOI:** 10.64898/2026.04.22.719835

**Authors:** Yorick van de Grift, Tamina Weiss, Guiliano Castellano, Wenjing Zhong, Łucja Biechońska, Kim Hellerstedt, Mikael Sigvardsson, Stefan Koch, Claudio Cantù

**Author notes:** Bluesky: @yorickgrift.bsky.social; @claudiocantu81.bsky.social (Yorick van de Grift, Claudio Cantù).

## Abstract

Wnt/β-catenin signalling drives gene expression in a plethora of processes during development and tissue homeostasis. Yet, little is known about how β-catenin, the transducer of the pathway, gives rise to context-specific transcriptional outcomes. For example, in the intestinal epithelium, stem cells and secretory cells lie adjacent to each other, are both exposed to and dependent on Wnt signals, but display divergent identities. We find that, in this tissue, GATA transcription factors functionally interact with Wnt/β-catenin signaling to drive differential cell identity. This occurs because GATA selectively repress Wnt target secretory-lineage genes in stem cells. Mechanistically, we found that GATA factors physically engage with the β-catenin transcriptional complex, and that this association is mediated by the bridge protein BCL9/9L. Our work implicates GATA factors as context dependent modulators of the Wnt/β-catenin transcription, shows that the GATA/BCL9 module acts as a switch at the crossroads of cell identity decision, and uncovers how this epithelium maintains the fine balance between self-renewal and differentiation.

## Introduction

The Wnt/β-catenin signaling pathway is a conserved cell-to-cell communication mechanism that coordinates a wide spectrum of processes by activating the expression of Wnt target genes^1^. Wnt/β-catenin assumes a crucial role in nearly all facets of embryonic development and adult homeostasis, spanning pluripotency, stemness, proliferation, and differentiation^2^. Its aberrant activation has been linked to many diseases such as developmental irregularities and various forms of cancer^2^. While much work has been dedicated to uncovering the principles of Wnt mediated control of stem cell identity and the activation of proproliferative transcriptional programs upon its dysregulation in cancer, Wnt/β-catenin signaling can also induce differentiation. However, how activation of this pathway can result in divergent cellular outcomes remains unclear.

WNT ligands activate an intracellular cascade that culminates in the accumulation of β-catenin and its translocation to the nucleus^3^. Nuclear β-catenin binds to TCF/LEF transcription factors positioned at Wnt Responsive Elements (WRE) on the chromatin together with a host of transcriptional cofactors that include BCL9, its paralog BCL9L, PYGO1/2^4–6^, the ChiLS (ChIP/LDB (LIM domain-binding protein) and a tetramer of SSDP (single-stranded DNA-binding protein)^7^ along with members of the SWI/SNF (BAF) complex and RNA polymerase II associated factors^8,9^. However, a model in which a universal β-catenin transcriptional complex drives the expression of Wnt target genes fails to explain a growing body of evidence showing that—beyond a common set of targets—Wnt activation induces tissue- and cell-type specific gene expression^10,11,12^. How does β-catenin transform the signals from the extracellular environment into the appropriate cell-specific gene expression program?

Here we address this question using the small intestinal epithelium as model system. Wnt/β-catenin drives both ISC stemness and secretory differentiation in the intestinal epithelium^13–15^. Turnover of this tissue relies on the Wnt-induced rapid division of intestinal stem cells (ISC), and their differentiation into secretory Paneth Cells (PC). Both cell types are strictly dependent on Wnt/β-catenin and, despite being physically adjacent at the bottom of the intestinal crypt, display two divergent cell identities^13–16^. This is an ideal model system to investigate how different competencieemerge despite in the presence of the same inductive signal (Fig 1A).

**Figure 1.**
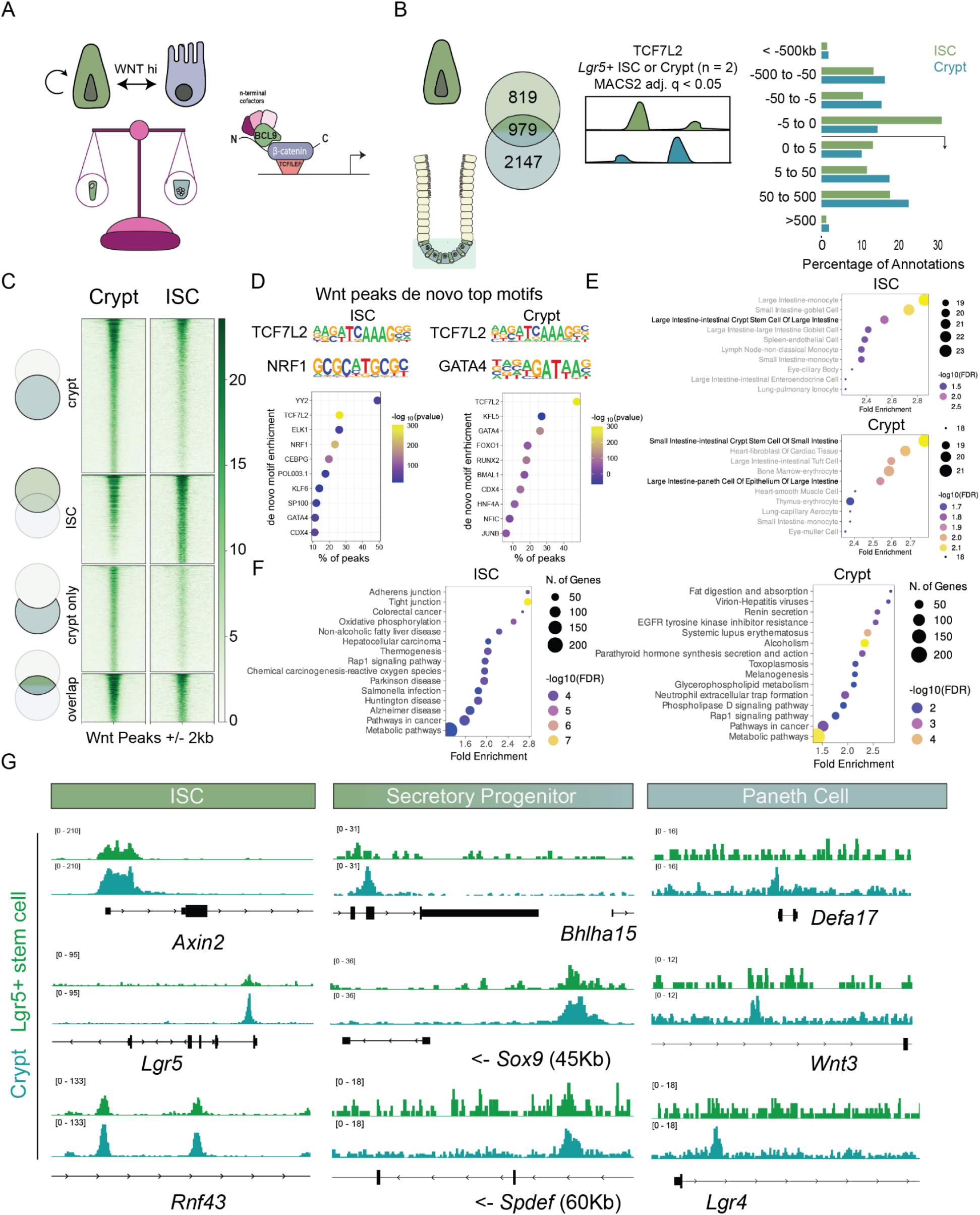
The genomic landscape of TCF7L2 binding in the intestinal crypts. A) Illustration of the balance between Wnt-mediated proliferation and differentiation in the intestinal crypt. B) Schematic representation of the experimental setup and overlap between ISC and Crypt datasets as displayed by a Venn diagram and genomic region annotation of TCF7L2 peaks by GREAT^20^. C) Signal intensity plots of TCF7L2 ChIP-seq signal within peak called regions of ISC, Crypt, Crypt only or overlap spanning the peak regions +/-2kb comparing ISC versus Crypt signal. D) De novo motif/consensus sequences discovered by HOMER^52^ in the TCF7L2 peaks [q(Benjamini–Hochberg False Discovery Rate adjustment) < 0.0001 for all motifs displayed]. The TCF7L2 peaks are statistically most enriched for their consensus TCF7L2 motif, but secondary motifs differ between ISC and Crypt peaks. E) Gene set enrichment analysis by ShinyGO^22^ of genes linked to peaks by GREAT to identify enrichment of markers associated with cell types in ISC versus unique Crypt peak sets. Marker genes were based on the Tabula Sapiens database^21^. F) Top 15 KEGG pathways enriched by ShinyGO (Fold enrichment and FDR < 0.05, background all protein coding genes) in ISC and unique Crypt peak associated gene sets. Dot size indicates the gene count, color indicates the negative decadic logarithm of the q-value after multiple testing correction (Benjamini-Hochberg False Discovery Rate adjustment). G) ISC, Secretory Progenitor and Paneth cell marker gene loci serve as examples of overlapping or unique normalized TCF7L2 ChIP-seq signal in ISCs versus Crypt (green and turquoise tracks, respectively). Green tracks represent TCF7L2 signal in ISC and turquoise tracks represent TCF7L2 signal in the Crypt.

We identify GATA transcription factors as a regulatory switch between the proliferative and the differentiation-inducing transcriptional programs mediated by Wnt/β-catenin signaling. In particular, GATA4 is expressed in ISCs, where it physically occupies a substantial subset of TCF7L2 (the main TCF/LEF protein in this tissue^14^) genomic binding sites linked to genes involved in secretory lineage differentiation. GATA4 knockout results in the upregulation of these genes, underlying its repressive nature, and accordingly in an increased number of secretory cells. Moreover, we show that modulation of Wnt/b-catenin signaling by GATA4 is direct, as evidenced by its ability to repress the SuperTOPFlash transcriptional reporter. Finally, we show that GATA colocalizes with β-catenin in nuclear foci, and that their association is dependent on the multifunctional bridge protein BCL9/9L. This work establishes GATA transcription factors as relevant tissue-specific players of Wnt/β-catenin signaling, and identifies a new mechanism that mediates the balance between the stem cell and secretory lineage identity in the intestinal epithelium.

## Results

### Genome-wide binding of TCF7L2 is rearranged during secretory differentiation

Previous work has shown that TCF7L2 is the dominant TCF/LEF in the intestinal epithelium^14^, a fact that we confirmed in intestinal organoids (Supplementary Fig 1A). We compared the TCF7L2 genome-wide binding by ChIP-seq in FACS sorted Lgr5^+^ intestinal stem cells (ISCs) versus isolated entire crypts^17^ (Fig 1B). Our rationale is that intestinal crypts contain both Wnt-responsive ISCs and Paneth cells (and other secretory progenitors), and this would allow us to discern Wnt/TCF7L2 targets in self-renewing versus differentiating cells. This approach yielded 1798 statistically significant peaks in ISCs and 3126 in the whole crypt (MACS2^18,19^ q <0.05). 69% of the crypt peaks (2147) were not present in ISCs, implying that during secretory differentiation the β-catenin transcription complex acquires novel binding sites (Fig 1B-C). This is also exemplified by distinct genomic distributions: in ISCs, TCF7L2 is mostly enriched at promoters, but more at distal regions in the other epithelial cells (Fig 1B).

*De novo* motif/consensus analysis of the DNA underlying the TCF7L2 binding sites in ISCs and crypts revealed the signature of a plethora of transcription factors. As expected, the top statistically enriched motif in both datasets is that of TCF7L2, but we observed differential enrichment of other transcription factor families, in particular GATA and KLF (Fig 1D).

We used GREAT^20^ to assign ISCs and crypt peaks to genes (Supplementary table 1). Gene set enrichment for cell markers based on the Tabula Sapiens database^21,20^ shows that genes associated to ISCs peaks were enriched for an ‘intestinal crypt stem cell’ expression program, whereas crypt peaks were linked to both ISCs and Paneth cell markers (Fig 1E) [Cell markers were considered significant with a q-value < 0.05 after multiple test correction (Benjamini-Hochberg False Discovery Rate adjustment)]. KEGG pathway analysis of peaks associated to ISCs and crypts genes yielded different biological processes consistent with their known cellular functions. In ISCs, for example, TCF7L2 was associated with genes related to cell proliferation and migration, including adherens and tight junctions, colorectal cancer and hepatocellular carcinoma. In crypts, TCF7L2 was linked to genes involved in metabolic processes such as fat digestion and absorption, and glycerophospholipid metabolism (Fig 1F). We consider these observations a powerful validation of our approach. The rearrangement of the genome-wide binding of TCF7L2 likely occurs gradually during secretory differentiation. TCF7L2 binding is already present in ISCs at known target genes of early secretory cells such as *Sox9* but absent at markers for mature Paneth cells like *Defa17* and it thus must be acquired at later stages of epithelial differentiation (Fig 1G).

### Global changes in chromatin accessibility underlying the TCF7L2 binding sites only partially follow secretory differentiation

We reasoned that gradual changes in chromatin accessibility could explain the different TCF7L2 binding patterns observed between ISCs and crypts. To test this, we analysed single cell-ATAC-seq from intestinal organoids^23^, computationally generated pseudobulk chromatin tracks for ISCs, secretory progenitor cells and Paneth cells (Fig 2A), and performed MACS3 peak calling on each set (q <0.05). Despite extensive changes in gene expression and phenotype, surprisingly little differences in chromatin accessibility occur during secretory differentiation: ISCs and Paneth cells share 69% of their accessible regions (Fig 2B, Supplementary figure 2B). The chromatin accessibility of TCF7L2 binding sites throughout secretory differentiation appears surprisingly static with only a fraction of regions changing their state from open to close or from close to open (Fig 2C). This observation extended to the TCF7L2-occupied binding regions. 49% of all TCF7L2 sites are accessible across all three cell states (Fig 2B, D). Only a small subset (6%) of the total TCF7L2 sites is closed in ISCs and gains chromatin accessibility during secretory differentiation. These regions are mostly associated with mature Paneth cell markers, such as *Defa17* (Fig 2F). Also, in ISCs only a relatively small subset of TCF7L2 binding sites is exclusively accessible in the stem cell population and includes ISC marker genes like *Ascl2* and *Kcnq1* (Fig 2F). However, many TCF7L2 binding sites that are linked to Wnt target genes such as *Lgr5* and *Spdef* do not change chromatin accessibility upon secretory differentiation (Fig 2F). We conclude that chromatin accessibility *per se* may be a poor predictor to understand the molecular cause underpinning the changes in the TCF7L2 genomic binding pattern observed during secretory differentiation. Interestingly however, *de novo* motif analysis of the DNA sequences underlying TCF7L2 binding sites in ISCs versus crypts uncovered a differential enrichment of a shortlist of potential cofactors at sites unique to differentiating cells (Fig 1D, Fig 2E). Among these, our attention was drawn again to GATA motifs (Fig 2E), as GATA transcription factors have been previously linked to proliferation and secretory differentiation in the intestine^24–31^.

**Figure 2.**
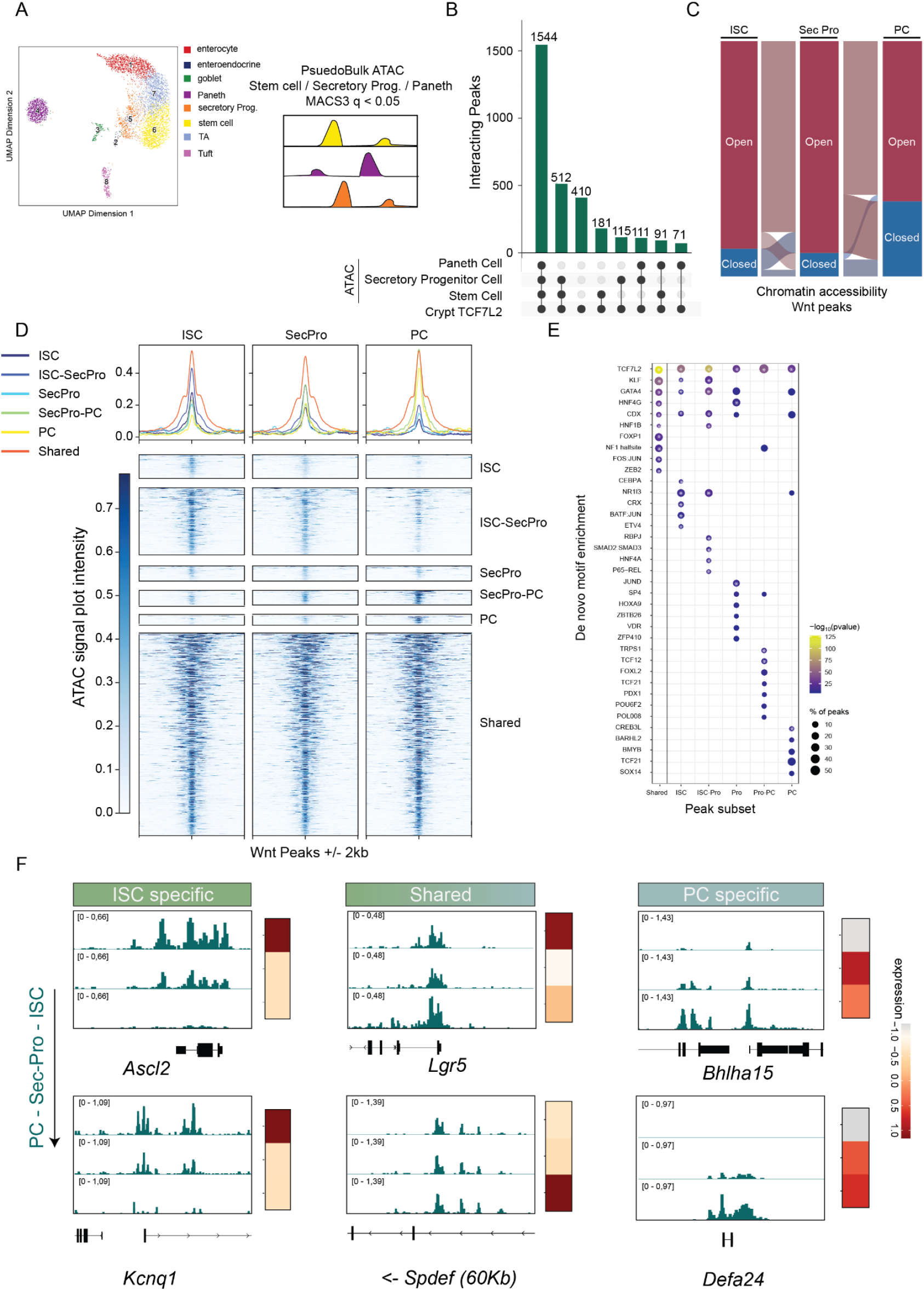
Chromatin accessibility underlying TCF7L2 binding sites. A) UMAP plot displaying scATAC-seq data from small intestinal organoids highlighting its respective cell types including ISCs, Secretory progenitors and Paneth cells that were used for further analysis. B) UpSet plot showing peak overlaps between peak called regions in pseudobulk ATAC-seq from ISCs, Secretory progenitors, Paneth cells and crypt TCF7L2 ChIP-seq. The highest overlap was between ATAC and TCF7L2 peak called regions was in regions that do not change accessibility between cell types. C) Alluvial plot that shows the ATAC-seq dynamics at TCF7L2 peaks. Only a minor set of regions undergo consecutive opening or closing during secretory differentiation, whereas most remain unchanged. D) Signal intensity plots of the pseudobulk ATAC signal of ISCs, Secretory progenitors and Paneth cells within the overlapping peak sets identified in B). The peak regions span the peak summit +/- 2kb. E) De novo motif/consensus sequences discovered by HOMER within the overlapping peak sets identified in B). All sets are statistically most enriched for the TCF7L2 consensus motif, but enrichments of secondary motifs differ between sets. Statistically significant motifs are highlighted by an asterisk [q(Benjamini–Hochberg False Discovery Rate adjustment) < 0.0001]. F) ISC, Secretory progenitor and Paneth cell marker gene loci serve as examples to highlight different temporal dynamics of pseudobulk ATAC signal across secretory differentiation and to what extend this correlate to gene expression in scRNA-seq in these respective subpopulations (heatmap to the right).

### GATA4 genome-wide binding overlaps with that of TCF7L2 on genes associated with secretory differentiation

Our data point to a model whereby expressed co-factors, including GATA4, the main GATA factor expressed in this context, might facilitate changes in Wnt target gene expression upon secretory differentiation. This model would predict the existence of a conspicuous overlap between GATA4 binding sites and Wnt target genomic regions. We tested this prediction by comparing TCF7L2 and GATA4 genome-wide binding via ChIP-seq^23^ (Fig 3A). Of 3126 statistically significant TCF7L2 peaks, 20% (623) colocalize with GATA4 (Fig 3A). This indicates that ca. one fifth of the Wnt/TCF7L2-dependent transcriptional outcome may be modulated by GATA4. GATA4 activity itself seems to be much broader, exemplified by a total of 15118 peaks (Fig 3A). We performed *de novo* motif search considering the GATA4/TCF7L2 overlapping sites. Here, both GATA4 and TCF7L2 motifs are more strongly co-enriched (Fig 3B). Of note, GATA motifs are positioned in the vicinity of TCF7L2 sites in a window that spans 200 nucleotides around the TCF7L2 binding sites (Fig 3C). We performed GREAT to link GATA4 and GATA4/TCF7L2 peaks to genes. The overall GATA4 peaks dataset was enriched in intestinal epithelial and enterocytic markers, including as *Alpi* and *Krt20* (Fig 3D-E). The GATA4/TCF7L2 peaks, on the other hand, are uniquely enriched for markers of Paneth cell identity, stem cells and early progenitors, such as, respectively, *Spdef, Lgr5* and *Sox9* (Fig 3D-E). Taken together, these data suggest that GATA4 associates to WREs found in regulatory regions of genes involved in cell lineage identity.

**Figure 3.**
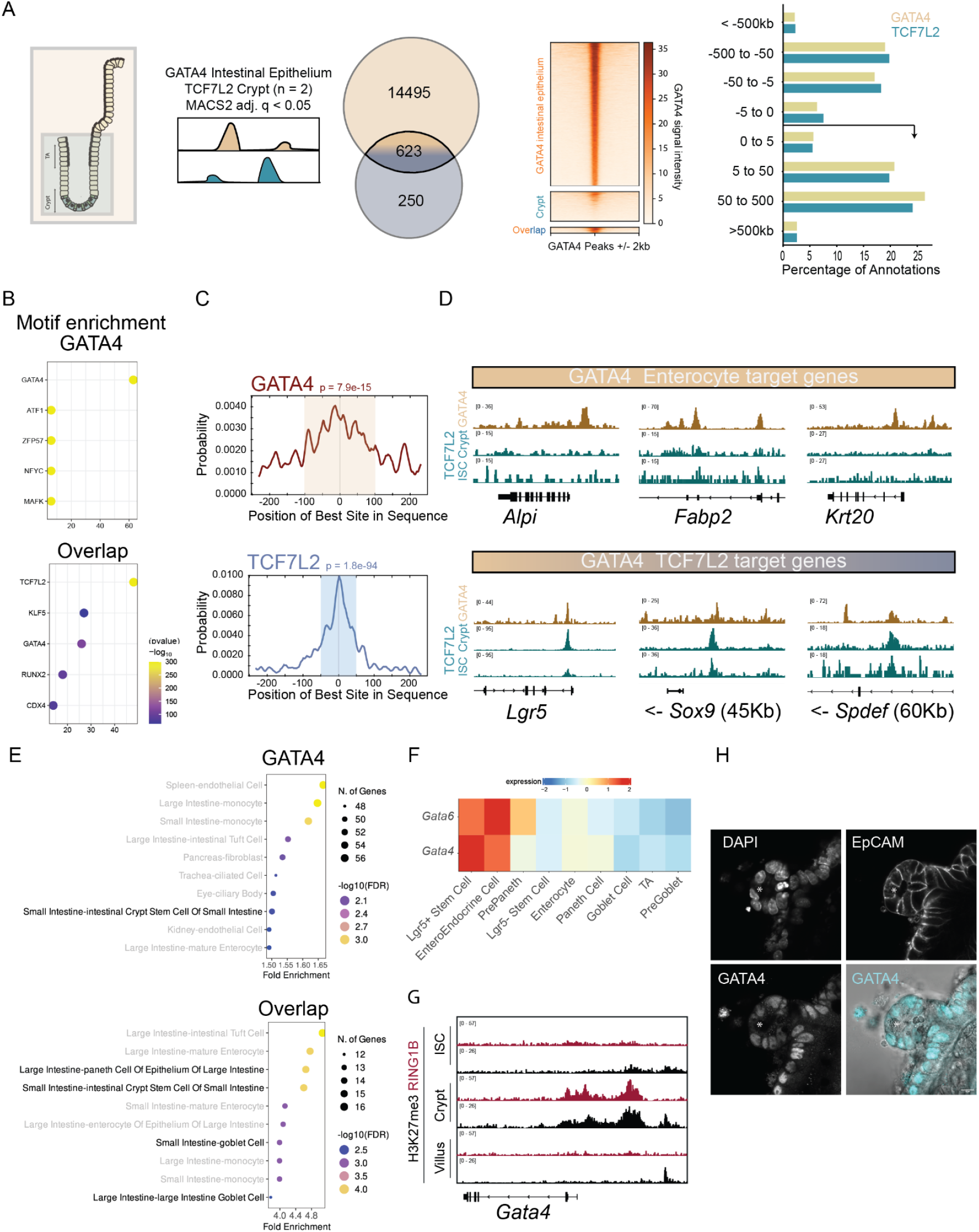
Figure Correlation between GATA4 and TCF7L2 in the intestinal epithelium. A) Illustration of the experimental setup and the overlap between GATA4 and Crypt datasets as displayed in a Venn diagram and signal intensity plot of the GATA4 ChIP signal within peak called regions of GATA4, Crypt TCF7L2 and overlap spanning the peak regions +/- 2kb. Comparative genomic region annotation of GATA4 and TCF7L2 peaks by GREAT is highlighted on the right. B) De novo motif/consensus sequences discovered by HOMER in the GATA4 peaks and overlapping peaks [q(Benjamini–Hochberg False Discovery Rate adjustment) < 0.0001 for all motifs displayed]. The GATA4 peaks are predominantly enriched for GATA4 motifs whereas the overlapping regions are strongly enriched for both TCF7L2 and GATA4 motifs. C) Centrality enrichment within overlapping peaks by Centrimo of GATA4 motifs (top) and TCF7L2 motifs (bottom). D) Enterocyte marker genes and TCF7L2 target gene loci server as examples of overlapping or unique GATA4 normalized ChIP-seq signal in Enterocyte versus Crypt and indicates GATA4’s dual role independent of TCF7L2 (orange and dark blue tracks respectively). Orange tracks represent the GATA4 signal and dark blue tracks represent normalized TCF7L2 signal in ISCs and the Crypt. E) Gene set enrichment analysis by ShinyGO of genes linked to the total set of GATA peaks or overlapping peaks by GREAT to identify enrichment of markers associated with cell types. Marker genes were based on the Tabula Sapiens database. F) Heatmap highlighting the relative gene expression of Gata4 and Gata6 expressing both genes in different cell types in intestinal organoids. G) H3K27me3 normalized ChIP-seq signal enrichment, a repressive histone mark, and Ring1b, a component of the Polycomb repressive complex 1, at the Gata4 promoter in ISCs, Crypt and Villi indicative of differential activation of Gata4 in each compartment. H) Representative confocal microscopy image of GATA4 staining in the crypt of small intestinal organoids. Nuclei are visualized with DAPI and EpCAM is used as a cell membrane marker. Each channel is separately displayed in grey scale and GATA4 signal in turquoise is overlayed to the brightfield image. Paneth cells are indicated by the asterisk.

### GATA4 acts in ISCs and early progenitors to repress secretory lineage genes

We have established that GATA4 and TCF7L2 both bind genomic regions in the intestinal epithelium that are linked to secretory lineage differentiation. However, from the binding profiles one cannot discern neither i) in which cell type GATA4 is active, nor ii) if it influences the transcription of Wnt/TCF7L2 target genes positively or negatively. Single cell transcriptomics profiling (Fig 3F) shows that *Gata4* (and the potentially redundant *Gata6*) is predominantly expressed in ISCs and enteroendocrine cells but gradually lost in other secretory lineages (Fig 3F). Consistently, the *Gata4* promoter is marked both by H3K27me3—a histone modification associated with repression— and by the Polycomb repressive complex 1, in crypt cells but not in ISCs nor along the villi (Fig 3G), indicating that its expression is specifically repressed in the secretory lineage. We confirmed this by immunofluorescence and observed that GATA4 protein is detectable along the crypts but specifically absent in Paneth cells (indicated by the asterisk) (Fig 3H).

The second point concerns the functional consequence of the GATA4/TCF7L2 interplay. Their co-occupancy is linked to genes involved in secretory differentiation, while *GATA4* expression is restricted to ISC and absent in secretory cells (Fig 3F-H). We reasoned that a way out of this apparent paradox is that GATA4 acts as a transcriptional repressor of Wnt/TCF7L2 secretory targets, thereby preventing ISC differentiation. This hypothesis is supported by several lines of evidence. First, *Gata4* knockout in the intestinal epithelium leads to an increase of active chromatin marks in the proximity of GATA4 targets^23^. Second other authors have shown that loss of *Gata4* in the mouse intestinal epithelium causes an increase in secretory cells^25–27^. Third, the consequence of the loss of *Gata4* (and similarly of *Gata6*) in the intestinal epithelium is reminiscent of specific Wnt phenotypes. For instance, earlier studies have shown that knockout of *Bcl9/9l* in addition to heterozygous mutations of *Ctnnb1* (encoding for β-catenin) caused ISCs exhaustion in favor of secretory hyperplasia pointing to BCL9/9L as another potential negative regulator of secretory differentiation^32^. Consistently, loss of *Gata4/6* and *Bcl9/9l* show a similar histological phenotype (compare Fig 7A in *Beuling E, Baffour-Awuah et al. 2012*^27^ with Supplementary figure 2 in *Deka J et al. 2010*^33^). The authors correctly concluded that *Bcl9/9l* loss leads to no gross malformations, further confirmed by the fact that the conditional knockout mice remained alive and well for the subsequent six months. However, our impression from their images is that they might have neglected a subtle increase of secretory cells (see supplementary figure 2 in *Deka J et al. 2010*^33^). To definitely test if loss of *Bcl9*/*9l* causes a compatible phenotype to the loss of *Gata4/6*, we generated *Rosa26*^*CreERT2*^*;Bcl9*^*fl/fl*^*;Bcl9L*^*fl/fl*^ small intestinal mouse organoids in which *Bcl9/9l* alleles can be conditionally deleted, and performed scRNA-seq at 72 hours after tamoxifen administration (Fig 4A). The results unambiguously showed an increase in secretory cells populations upon *Bcl9/9l* loss (Fig 4B). A random permutation test confirmed that Goblet and Paneth cells grew significantly in number (FDR < 0.05, abs(Log2FD) > 0.58) (Fig 4C). We carefully quantified this effect independently by RT-qPCR. Loss of *Bcl9/9l* resulted in a 2-fold increase of Paneth cell marker genes *Lyz1* and *Wnt3*, whereas ISCs and Wnt target genes *Axin2* and *Lgr5* remain unaffected (Fig 4D). Our experiments demonstrated that loss of *Gata4/6* or *Bcl9/9l* in the intestinal epithelium results in matching phenotypes.

**Figure 4.**
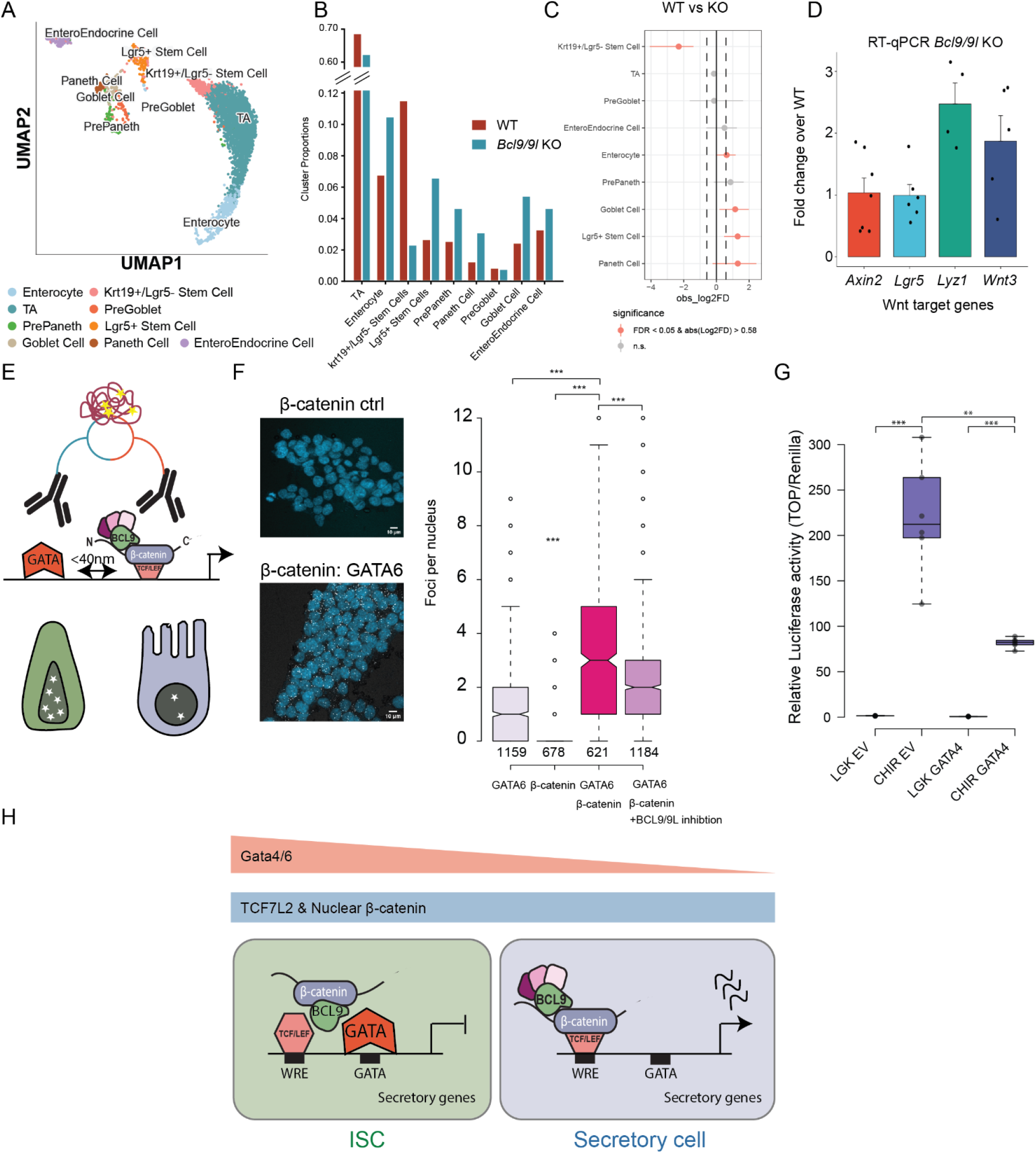
GATA4 engages the β-catenin transcriptional complex via BCL9/9L. **A)** UMAP plot displaying scRNA-seq data from small intestinal organoids highlighting its respective cell types. B) Barplot that visualizes the respective changes in cluster proportions 72h hours post Bcl9/9l deletion versus wildtype organoids. Cluster sizes are displayed in proportion to the total number of cells. C) Statistical significance of observations in B). A permutation test is used to calculate a p-value for each cluster, and a confidence interval for the magnitude difference is returned via bootstrapping (FDR < 0.05, abs(Log2FD) > 0.58). D) Gene expression measurement by RT-qPCR 72h post Bcl9/9l knockout of stem cell marker genes: Axin2 and Lgr5 and Paneth cell marker genes: Lyz1 and Wnt3 showing that Bcl9/9l knockout only affects gene expression of Paneth cell maker genes. Gene expression is displayed as relative fold change to the wildtype. Rpl13a was used as a reference gene. Error bars show mean ± SD among independent experiments (n = 2-3). E) Schematic illustration of proximity ligation assay. F) Representative confocal microscopy images from the proximity ligation assay showing GATA6 only or GATA6 + β-catenin (right). Quantification of proximity ligation assay signal, in foci per nucleus (left). GATA6 + β-catenin has significantly more interaction than the controls (p < 0.001) and pretreatment with C1, a BCL9/9L-β-catenin inhibitor, significantly reduces this interaction. Counted nuclei = 3642. G) TOPFlash assay investigating the effect of GATA4 on the activation of Wnt/β-catenin signaling. HEK293T cultured in 10 nM LGK, were transfected with a control empty vector (EV) or pXJ-myc-GATA4 with or without 1 μM CHIR99021^53^. All cells were transfected with the TCF/LEF-luciferase reporter SuperTOPFlash and Renilla control plasmids^37^. WNT-OFF and WNT-ON conditions were tested with 3 replicates per condition. Activation of Wnt/b-catenin signaling was measured as relative luciferase activity. Data was visualized as box and whiskers plots by BoxPlotR^54^. A Student’s t test was used for pairwise comparison of groups, and significance values (p-value) are displayed as asterisks above the boxes. A p-value < 0.05 was used to determine statistical significance. H) Schematic illustration of the final model.

### GATA physically associates with the β-catenin transcriptional complex through BCL9/9L

We asked whether GATA4 may exert its control of secretory lineage by physical interaction with the β-catenin transcriptional complex. The overlapping genomic signal between GATA4 and TCF7L2 were suggestive of such possibility. However, while ChIP-seq identifies regions bound by these factors in independent experiments, it does not allow to distinguish if they co-occupy the same locus simultaneously. We applied CentriMO^34^ to investigate the positional distribution of GATA4 motifs relative to the peak center of TCF7L2 at GATA4/TCF7L2 co-occupying regions in the intestinal crypt. TCF7L2 motifs are centrally enriched as expected (1.8E-94). GATA4 motifs are not as locally enriched and are more broadly distributed compared to the TCF7L2 peak center (7.9E-15) (Fig 3C), indicating that TCF7L2 and GATA4 binding might not be mutually exclusive. To test the vicinity of GATA with the active Wnt/b-catenin transcriptional complex, we performed proximity ligation assay (PLA), which uses microscopy to detect a signal emerging when two proteins of interest are in close physical proximity (within 40 nm)^35^, in HCT116 cells where Wnt/b-catenin signaling is constitutively active (Fig 4E). While GATA4 is predominantly expressed in the small intestine, GATA6 is the main GATA transcription factor present in the colon epithelium. As HCT116 cells are derived from the colon, we tested GATA6 colocalization with b-catenin. Foci of β-catenin plus GATA6 are detected proximally to the chromatin and significantly enriched over the control (p < 0.0001, median 3 foci per nucleus) (Fig 4F).

The similarity of in vivo phenotypes indicates that the bridge protein BCL9/9L may have a role in the GATA6/b-catenin observed proximity. To test this, we treated HCT116 cells with a recently developed inhibitor of the BCL9/9L-β-catenin interaction (C-1)^35^. C-1 administration strongly reduced the number of foci GATA-β-catenin detected by PLA (p < 0.001) (Fig 4F). We conclude that GATA transcription factors and β-catenin physically engage at the chromatin of the intestinal epithelium through BCL9/9L. Our model would imply that GATA has a repressive effect on Wnt/b-catenin signalling and that this may be mediated by its interaction with the Wnt-dependent complex via BCL9/9L, possibly even independently by the presence of GATA motifs. We tested this by employing the widely used transcriptional SuperTOPFlash reporter that specifically allows to measure Wnt/β-catenin exclusive transcriptional activation^37^. Overexpression of GATA4 significantly abrogated the transcription from SuperTOPFlash upon strong Wnt/β-catenin pathway activation achieved using CHIR99021 (p < 0.01) (Fig 4G). Collectively, our findings show that GATA4/6 can act as a transcriptional repressor of Wnt-dependent transcription by directly interacting with the β-catenin transcriptional complex through BCL9/9L and that this is required to finetune Wnt/b-catenin signalling and repress secretory differentiation.

## Discussion

A model in which a universal β-catenin transcriptional complex drives the expression of Wnt target genes fails to explain a growing body of evidence indicating that Wnt activation induces tissue- and cell-type specific gene expression^10,11^. Given the many roles in diverse cellular contexts attributed to this pathway, one key question remains unanswered: how does β-catenin mechanistically transduce the extracellular signal into appropriate cell-specific gene expression programs? One explanation entails tissue-specific transcription factors that help coordinate Wnt/β-catenin controlled gene expression. This includes lineage-specific partners of β-catenin that either act competitively, like SOX transcription factors^38,39^ and OCT4^40^, or synergistically such as TBX3^40,41^ and c-JUN^42^.

Here we find an additional instance supporting this interpretation. We tackled the intestinal crypt, and difficult case where ISCs and Paneth cells, two Wnt/β-catenin-dependent cell types lie physically adjacent but display two divergent cell identities in response to Wnt. ISCs and Paneth cells are exposed to the same WNT ligands^43^, equally express TCF/LEF transcription factors^14^ (supplementary figure 1A) and display no discernible difference between nuclear β-catenin levels^44^. However, these cell types activate different Wnt-dependent gene expression programs. Here we identify a new mechanism that explains how.

We detected a genome-wide repositioning of the Wnt/β-catenin transcriptional complex, that rewires from ISCs to differentiating cells to control genes involved in the secretory lineage. This genome-wide shift is not globally driven by major changes in chromatin accessibility at WREs and other regulatory regions. Instead, we uncover that GATA4 and 6 act as transcriptional co-factors of the β-catenin transcriptional complex in a BCL9/9L dependent manner, a protein known to finetune Wnt/β-catenin target gene expression in this tissue^32^.

GATA4/6 have been implicated in various biological processes in the intestinal epithelium, including proximal-distal patterning of the enterocyte lineage, crypt cell proliferation and secretory differentiation^24–29^. But GATA4/6 expression is high in ISCs and lost during secretory differentiation. This apparent paradox is resolved by showing that these proteins act as transcriptional repressors on secretory lineage genes—that are also Wnt targets—in ISCs. Our observation is further corroborated by conditional knockout studies that show that a loss of *Gata4/6* in the mouse results in the increase of secretory markers, including the lineage specificator and Wnt target gene *Spdef*. Combined *Gata4/6* and *Spdef* loss partly abrogates the secretory hyperplasia—thereby rescuing the effect of a loss of *Gata4/6* alone—underscoring the role of GATA4/6 as secretory-lineage repressors^26^.

Previous reports show that histone methyltransferase MLL1 is an upstream positive regulator of *Gata4/6* expression in ISCs^46,45^. Its primary function is to prevent Polycomb mediated repression of ISC gene expression by actively replacing the repressive H3K27me3 mark with H3K4me3, which is needed to maintain *Gata4/6* expression and prevent secretory hyperplasia. We confirmed that, indeed, H3K27me3 marks are absent on the *Gata4* promoter in ISCs and that deposition of this repressive mark and the binding of the Polycomb repressive complex 1 correlates with loss of *Gata4* expression itself (Fig 3G). We expand on this regulatory axis in ISCs by highlighting a novel molecular mechanism by which GATA4/6 control and finetune gene expression to modulate crypt cell proliferation and secretory differentiation.

Our data are compatible with a model in which GATA4/6 associates with β-catenin through BCL9/9L in ISCs. In the permissive environment of the ISC niche, where Wnt/β-catenin signalling is activated and differentiation programs are repressed, GATA4/6 has a repressive effect that finetunes Wnt/b-catenin signalling and prevents activation of Wnt-dependent secretory differentiation. (Fig 4H). This model is supported by evidence of a similar function in the adult heart, breast cancer and hepatocellular carcinoma^47,48,49^, which indicates that this mechanism is not restricted to the intestine but more universally adopted in GATA4/6 dependent tissues. We expand this concept by showing that BCL9/9L is required for GATA4/6 mediated finetuning of Wnt/β-catenin target gene expression.

Intriguingly, inspection of published data on protein-protein interactions in colon cancer cells revealed that the interaction between GATA6 and β-catenin had already been detected in this context, and that these cancer cells are highly co-dependent on GATA6 for their proliferation^50^. This finding is corroborated in *Apc* knockout colon organoids, where additional loss of *Gata6* supresses tumorigenesis and the organoids revert to a crypt-like structure^51^.

We propose that GATA4/6 act as gatekeepers between Wnt/β-catenin-mediated proliferation versus differentiation. This not only represents the explanation of how two adjacent cells, exposed to the same signal, can transduce it in different manners, but also offers a tantalizing therapeutic avenue that will exploit the sensitive dependence that colon cancer stem cells display on GATA proteins for their proliferation

## Acknowledgments

We thank all the members of the Cantù lab for the continues scientific input. The computations and data handling were enabled by resources provided by the National Supercomputer Centre (NSC), funded by Linköping University. We thank Peter Münger at the NSC for assistance concerning technical and implementational aspects in making the codes run on the Sigma resource.

## Author contributions

Conceptualization: Y.v.d.G, C.C.; Methodology: Y.v.d.G, T.W., G.C., W.Z., L.B., K.H.; Data curation: Y.v.d.G., C.C.; Formal analysis: Y.v.d.G.; Investigation: Y.v.d.G., T.W., G.C., W.Z., L.B., K.H.; Funding: C.C., S.K., M.S.; Project administration: C.C.; Visualization: Y.v.d.G.; Writing – original draft: Y.v.d.G., C.C.; Writing – review & editing: Y.v.d.G., T.W., C.C.

## Data availability

All study data are included in the in the article and/or supplementary information. Raw data from previously published work was processed and reanalysed for this study (Supplementary table 4). All data generated in this studyis deposited on Array-Express and can be accessed at E-MTAB-16966.

## Methods

### Mouse husbandry

Animal housing and experimentations were performed according to the Swedish laws and guidelines under the ethical animal work license obtained by C.C. at Jordbruksverket (Dnr 2456-2019). Animals were kept in Allentown NexGen IVCs, floor area 500cm^2^. The maximum number of mice was four per cage. Cages were supplied with aspen wood shavings as bedding and two types of shredded paper as nesting material. Paper tubes were provided as additional enrichment. Temperature was set at 21±2°C, humidity to 45-65% and light cycle to 12h/12h (7.00am/7.00pm). The animals had unrestricted access to sterilized drinking water, and *ad libitum* access to a pelleted and extruded mouse diet in the food hopper. Mice were housed in a barrier-protected specific pathogen-free unit. The specific pathogen-free status of the animals was monitored frequently and confirmed according to FELASA guidelines by a sentinel program. The mice were free of all viral, bacterial and parasitic pathogens listed in FELASA recommendations^56^. *Bcl9*^*fl/fl*^*;Bcl9l*^*fl/fl*^*;Rosa26*^*CreERT2(HET)*^ mice backcrossed to JAX Swiss Outbred mice (strain 034608) were used to generate small intestinal organoids.

### Isolation and culture of mouse small intestinal organoids

Mouse small intestinal organoids were established as previously described^57^. Crypts were isolated from the entire length of the small intestine of one *Bcl9*^*fl/fl*^*;Bcl9l*^*fl/fl*^*;Rosa26*^*CreERT2*(HET)^ animal to establish the organoid line. Organoids were culture in 15ul BME (3434-005-02, Trevigen) droplets in culture medium containing advanced DMEM/F12 (12634010, ThermoFisher Scientific) supplemented with 100U/ml Penicillin/Streptomycin (15276355, ThermoFisher Scientific), 2mM Glutamax (11574466, ThermoFisher Scientific), 10mM HEPES (15630-056, ThermoFisher Scientific), and 1.25mM N-acetylCysteine (A9165-5G, Sigma Aldrich), freshly added mEGF (50ng/ml, 315-09, PeproTech), 2% Noggin-Fc conditioned media (N002-100ml, AVSbio), 2% Rspo3-Fc conditioned media (R001 – 100ml, AVSbio) at 37°C and 5% CO_2_. For passaging, cell culture medium was removed and BME was broken into small pieces by scraping, followed by vigorous pipetting with ice-cold advanced DMEM/F12. Crypts were centrifuged at 400g for 5 min at 4°C. The supernatant was carefully removed, the pellet was resuspended in 50:50 advanced DMEM/F12:BME and plated on pre-warmed culture plates. Medium was refreshed every other day, and organoids were split once a week in a 1:3 ratio. Organoids were grown untill day 5 and treated with 1μM 4-OHT (H7904-5MG, Sigma-Aldrich) for 24 hours to delete *Bcl9/9l*. Organoids were harvest at day 8 for qPCR or scRNA-seq.

### RNA isolation, cDNA synthesis an qRT-PCR

Total RNA was isolated from small intestinal organoids using QIAzol (79306, Qiagen) according to the manufacturer’s guidelines, with the addition of 20ug glycogen during RNA precipitation to enhance RNA isolation. The RNA concentration was determined using a Nanodrop spectrophotometer. cDNA was synthesized from up to 2ug RNA using High-capacity cDNA Reverse Transcriptase kit (43-688-13, ThermoFisher Scientific) according to the manufacturer’s instructions. The resulting cDNA was diluted 10-fold for subsequent qRT-PCR analysis. qRT-PCR was performed using a Bio-Rad CFX96 Real-Time system. PCR reactions (total volume 20ul) were set up containing 10ul Ssoadvanced Universal SYBR (1725274, Bio-Rad), 0.5μM of each specific forward and reverse primer (10μM stock) and 5ul of diluted cDNA template. The reactions were set up in technical duplicates in 96-well qPCR plates. One negative control (no-RT) reaction was included for each sample/primer combination. Thermal cycling was performed according to the following settings: 10 min 50°C, 5 min 95°C, [10 sec 95°C, 30 sec 60°C]^40x^, 10 sec 95°C, 5 sec 65°C, 30 sec 95°C. Each run was completed with a melting curve analysis. Primer sequences are detailed in Supplementary Table 4.

### Cell culture

Human HCT116 colon cancer cells (a kind gift from Stefan Koch) and HEK293T human embryonic kidney cells were routinely cultured in Dulbecco’s Modified Eagle Medium (DMEM) (41965062, ThermoFisher Scientific), supplemented with 100U/ml Penicillin/Streptomycin (15276355, ThermoFisher Scientific) and 10% Fetal Bovine Serum (FBS) (HCT116) (26140079, Thermo Scientific) or 10% Bovine Calf Serum (BCS) (HEK293T) (12133C-500ML, Sigma-Aldrich). Cells were split 1:10 twice a week.

### Microscopy

Images of small intestinal organoids were captured using confocal microscopy on a Leica Stellaris 5 with the LasX software. For imaging, the samples were cultured for 7 days and imaged on glass chamber slides (Ibitreat 80806, Ibidi). Intestinal organoids were fixed with 4% PFA (15670799, Fisher Scientific) for 15 min at room temperature, the reaction was quenched with 0.15M Glycine (04808822, MP Biomedicals) and the samples were permeabilized with 0.5% TritonX-100 (93443-100ML, Sigma-Aldrich) for 10 min at room temperature. The samples were blocked with 5% BSA (A9647-50G, Sigmal-Aldrich) solution for 2 hours. To visualize GATA4, the samples were stained with 1:200 primary antibody anti-GATA4 (ABIN6140975, Antibodies-online) overnight at 4°C, and 1:1000 secondary antibody Goat anti Rabbit Alexa Fluor 488 (A-21206, ThermoFisher Scientific) for 1 hour at room temperature. The organoids were incubated with 1 μM DAPI (D1306, Thermo Scientific) and 1:1000 EPCAM-APC Primary antibody (17-5791-82, ThermoFisher Scientific) for 10 min at room temperature. Fluorophores were excited as follows: DAPI at 405 nm, Alexa Fluor x/anti-GATA4 at 488 nm and EPCAM-APC at 633 nm. All images were acquired using a 25x water long-working distance objective. Fluorescence microscopy images were processed in Fiji using the Image 5D plugin^58^.

### Luciferase Assay

To conduct reporter assays, HEK293T cells were seeded in a 96-well plate overnight and co-transfected with 50ng firefly reporter, 5ng of renilla control, and 10ng of the plasmid of interest in each well. M50 Super 8x TOPFlash (SuperTOPFlash is found at Addgene, Plasmid #12456), M51 Super 8x FOPFlash (Addgene #12457) and pXJ-myc-GATA4 (Addgene #232647) as used before^59,60^. In the indicated samples cells were treated with 10nm LGK (S7143, Selleck Chemicals) (WNT-OFF) or 1μM CHIR99021 (SML1046, Sigma Aldrich) 6h after transfection. The dual luciferase assay was performed with some modifications as previously described^61^. After overnight incubation, cells were lysed in passive lysis buffer (25mM Tris, 2mM DTT, 2mM EDTA, 10% (v/v) glycerol, 1% (v/v) TritonX-100, pH 7.8) and agitated for 10 min. The lysates were then transferred to a flat-bottomed 96-well luminescence assay plate. Firefly luciferase buffer (200μM D-luciferin in 200mM Tris-HCl, 15mM MgSO4, 100μM EDTA, 1mM ATP, 25mM DTT, pH 8.0) was added to each well and the plate was incubated for 2 min at room temperature. The luciferase activity was measured using a SpectraMax iD3: Multi-Mode Microplate Reader (Molecular Devices). Subsequently, Renilla luciferase buffer (4μM coelenterazine-h in 500mM NaCl, 500nM Na2SO4, 10mM NaOAc, 15mM EDTA, 25mM sodium pyrophosphate, 50μM phenyl-benzothiazole, pH 5.0) was added to the plate, and luminescence was immediately measured. The data was normalized to the Renilla control values, performed in triplicate, and the Top/Renilla ratio was used as an indicator of β-catenin-driven transcription. Statistical analysis was performed in excel. Data was visualized in box and whiskers plots using BoxPlotR^54^. Student’s t-test was used to analyze the pairwise differences between groups and p < 0.05 was considered statistically significant. All experiments were performed in duplicate.

### Proximity Ligation Assay

Proximity ligation assay was performed using the Navinci Flex Cell Red kit (60025, Navinci) according to the manufacturer’s guidelines. If applicable HCT116 cells were pretreated with 10μM C-1 BCL9/9L inhibitor (DC70375, DC Chemicals) 24 hours prefixation. Cells were grown on a glass slide and fixed with 4% PFA for 15min at room temperature, the reaction was quenched with 0.15M Glycine and the samples were permeabilized with 0.5% TritonX-100 for 10min. The samples were blocked with 5% BSA solution for 2 hours. Antigens were detected using mouse anti-β-catenin (BD labs, 610154) and rabbit anti-GATA6 (ABIN6257236, Antibodies-online) at 1:200 dilutions. The assay was performed in duplicate. Images were acquired using a Leica Stellaris 5 confocal microscope with the LasX software imaging nuclei and PLA foci in separate channels. Acquisition settings were optimised on the controls and remained the same for all samples. Images were taken with a 25x water long working distance objective.

Image processing was performed using CellPose-SAM^62^ and Fiji^58^. For visualisation, channels were merged and assigned a colour using the plugin Image5D and scalebars were added. For quantification, channels were split in ImageJ and saved separately. The DAPI channel was used in CellPose-SAM. The default CPSAM model was used to segment nuclei based on the DAPI signal and acquire ROIs. ROIs were exported to the native ImageJ ROI format. The PLA foci channel images were loaded in Fiji and converted to a grey scale 16-bit image. Thresholding was applied equally to all images to create binary images. Open the set of ROIs from CellPose-SAM using the ROI manager. Before analysing the PLA foci include the ‘bounding rectangle’ and ‘limit to threshold’ options in set measurements. Select the first of the loaded ROIs, analyze particles and adjust the size to ‘0.0003-infinity’. Run the following macro to automatically count the number of PLA foci per nucleus ROI in each image. Measurements can be lifted to excel for further downstream analysis. Between 621 and 1184 nuclei were counted per condition, for a total of 3642 nuclei. Statistical testing was done with two-tailed *t*-tests in excel. Data was visualized using BoxPlotR^54^.

Macro code for automatic counting of foci:

c= roiManager(“count”);

for (i = 0; i < c; i++)
{ roiManager(“Select”, i);

run(“Analyze Particles…”, “show=Nothing display exclude include summarize add composite”);

}

roiManager(“show all without labels”);

### Chip-seq analysis

Raw fastq files from GEO: GSE65322, GSE68957, GSE71713 and GSE164940 were processed in the following manner. Reads were trimmed with bbmap bbduk (version 38.18)^63^ to remove adapter sequences using the following parameters: qtrim = r, minlen = 25 and trimq = 10. Alignment to the mm10 genome qas performed using Bowtie2 (version 2.2.5)^64^. Samtools (version 1.9)^65^ was used for BAM file creation, sorting, merging subsampling and indexing. Bam files were filtered to remove blacklist regions^66^. BigWigs were created withDeepTools (version 3.5.4)^67^ using BamCoverage with the -RPGC option for normalisation. Peaks were called using MACS2 (version 2.2.6)^18,19^ on a merge of two biological replicates against the input control, using the options -f BAM, -g mm on narrow-Peak mode with a q-value threshold of <0.05. Peak overlaps were performed using BEDtools (version 2.26.0)^68^.

For graph and visualisation purposes, replicate BAM files were merged with SAMtools into a single file. DeepTools was used to convert BAM files to normalized bigwig files (bamCoverage using -RPGC option), signal intensity plots, and profiles (computeMatrix reference-point followed by plotHeatmap). BigWigs were visualised using the IGV browser^69^. Peak annotation was done with GREAT (version 4.04) on default settings. Motif analysis were performed with HOMER (version 4.11) ^51^ using FindMotifsGenome.pl on default settings. SR Plot^70^ was used to visualize motif enrichment. Venn diagrams and Upset plots were created using Intervene (version 0.6.5)^71^. Gene ontology analysis was performed and plotted using ShinyGO (version 0.85.1) using the following options: species: human, pathway database: Co-expression.Tabula:Sapiens and KEGG, FDR cutoff: 0.05, pathways to show: 10, pathway size min:2 and pathway size max; 5000. Peak annotation by GREAT was used gene ontology analysis. Centrimo, part of the MEME-ChIP Suite^72^, was run on nonlocal mode. To run Centrimo, peak size was equalized to 500bp using Bedtools in the following manner: Peaks summits as defined by MACS2 were extended 250bp to either side with the bedtools slop function.

### scATAC-seq quality control

Mm10 aligned fragment files from GEO: GSE221421 were processed with ArchR^73^. Fragments were read into ArchR and a per cell quality control was performed on three characteristics of the data: 1) the number of unique nuclear fragments, 2) signal-to-background ratio and 3) the fragment size distribution. Default settings of a TSS enrichment score of >4 and >1000 unique nuclear fragments were used to exclude low quality cells from further analysis. Doublet enrichment scores were subsequently calculated for cellular barcodes using ArchR’s addDoubletScores() function on default settings (k = 10, knnMethod = ‘UMAP’ and LSIMethod = 1). Cellular barcodes with a doublet enrichment >1 were marked as potential doublets and removed using the filterDoublets() function.

### scATAC-seq processing

We used ArchR to construct an initial feature matrix of 500bp genomic tiles across all cells. To reduce the dimensions of the genomic tiles features, we used the iterative latent semantic indexing (LSI) in ArchR with the addIterativeLSI() function on default setting except for the option dimsToUse which was set to 2:50. Briefly, this procedure performs term infrequency-inverse document freuquency (TF-IDF) normalisation to upweight more informative features followed by an initial LSI reduction on the top accessible tiles followed by graph-based Louvain clustering to identify low resolution clusters in which feature counts are summed across all cells in each cluster to identify the top 25000 most variable features. This process was iterated twice by using the top 25000 features in the second iteration. The addCluster() function, which implements Seurat’s FindClusters() function, was used to identify clusters and dimensionality reduction was performed by UMAP embedding using the addUMAP() function. We used MAGIC^74^ to impute gene scores by smoothing signal across cells via the addImputeWeight() function. UMAP plots were generated using the plotEmbedding() function. Gene activity scores were inferred in scATAC-seq using ArchR’s addGeneIntergrationMatrix() function on default settings. scATAC clusters were annotated by integrating scRNA-seq and transferring cluster identity. To integrate scATAC with scRNA-seq, we used the addGeneIntegrationMatrix() function on default settings. As input, we used matching scRNA-seq from the same source (GSE221421) re-analysed in Seurat^75^ and converted to a singlecellExperiment object using the as.SingleCellExperiment() function from Seurat. The integration works by directly aligning cell from scATAC-seq with cells from the scRNA-seq by comparing the scATAC-seq gene score matrix with the scRNA-seq gene integration matrix via the FindTransferAnchors() function from Seurat. After receiving a cluster label, pseudo-bulk replicates were generated for each cluster using the AddGroupCoverages() function on default settings. The getGroupBW() function was used to export the pseudobulk ATAC signal per cluster as a bigWig file for further visualization. Peak calling was performed within each cluster using MACS3 with a threshold of q < 0.05 run through the addReproduciblePeakSet() function. Overlap between peak sets, Venn diagrams and Upset plots were created using Intervene. DeepTools was used generate signal intensity plots, and profiles (computeMatrix reference-point followed by plotHeatmap). Motif analysis were performed with HOMER using FindMotifsGenome.pl on default settings. SR Plot was used to visualize motif enrichment and generate the alluvial plot.

### scRNA-seq

Small intestinal organoids were harvested with ice-cold medium and transferred to a 15ml tube. Ice cold Advanced DMEM/F12 was added till 10ml, and the organoids were pelleted at 4°C for 5min at 400g. ‘Matrigel cloud’ was carefully removed and organoids were pelleted again in 10ml ice cold advanced DMEM/F12. Organoids were resuspended in 1ml TrypLE express enzyme, Phenol Red (12605010, Thermo Scientific) and incubated in a 37°C waterbath till the organoids were fully dissociated to single cells while gently shaking and occasional pipetting with a p1000 to prevent clumping. Cells were checked under a brighfield microscope every 5 min to assess dissociation. Once the organoids were fully dissociated to single cells the digestion was stopped with 500ul BCS, immediately topped till 10ml with ice cold Advanced DMEM/F12 and spin at 4°C for 5 min at 500g. The cells recovered from each condition were counted (range 100-200k) and scored for successful dissociation to single cells. The recovered cells were fixed according to the manufacturer’s procotocol (ECW02050, Parse BioSciences) including 0.5% BSA in the fixation solution (15260037, Fisher Scientific) and stored at -80°C. Before barcoding, cells suspensions were thawed and recounted and loaded on the first barcoding plate according to the manufacturer’s WT mega v2 sample loading table aiming at ∼10k cells per condition. Further barcoding and sequencing library generation was performed according to the manufacturer’s protocol, generating 6 sub-libraries, with a target of 20000 cells each (note that more experiments were mixed in the sequencing library, amounting to ∼1500 cells / condition / library). The final sub-libraries were evaluated on a TapeStation (Agilant) for size distribution and quantitative real time PCR (qPCR). 1 sublibrary was sequenced at a depth of 20.000 reads / cell on an Illumina NovaSeq 6000 S4 flow-cell with PE150 according to the results from library quality control and expected data volume by Novogene.

### scRNA-seq analysis

If applicable output fastq files were concatenated to a single file for each read end. Fastq.gz files per sublibrary were demultiplexed and aligned to hg38 using the Trailmaker pipeline by Parse Biosciences. Trailmaker was used to demultiplex the library. The unfiltered count matrix was used for further processing in the Trailmaker pipeline. The data was filtered based on cell size distribution, mitochondrial content, number of genes versus transcripts per cell, and filtered for doublets using automatic filter settings of the Trailmaker pipeline. Withing the Trailmaker pipeline, log-normalization, scaling and batch correction was performed with Seurat v4 and variance stabilizing transformation was performed with SCTransform v1 without removing covariates like mitochondrial, ribosomal or cell cycle genes. The top 2000 highly variable genes were used for integration with 30 principal components using the CCA method. The processed data was exported as a Seurat file and cell cluster identities were assigned in Seurat according to the markers in supplementary figure 1C (derived from PangloaDB^55^). ShinyCellR^76^ was used to plot the UMAP. Cluster proportions in WT versus BCL9/9L knockout were calculated in R as the relative proportion of each cluster to the total number of cells per condition and visualized with SRplot. Statistical significance of changes in cluster sizes between conditions were calculated and plotted using a random permutation test (FDR < 0.05) using the scProportionTest package^77^. Raw count matrix files from scRNA-seq matched to the scATAC-seq data from public repository GEO: GSE221421 were further processed in Seurat according to their standard pipeline. Briefly, data was filtered and subsequently normalised with nFeature_RNA values set to >200 and <4500 and percent. Mt values set to <10. For integration, datasets were anchored together with 3,000 integration features and 30 dimensions for identifying anchors before being clustered in accordance with the standard Seurat pipeline. Clusters were assigned a cell type based on known markers. This processed data was used to later assign cell types to scATAC-seq clusters.

## Additional Files

**Supplementary table 1**. Genomic coordinates of peak-called regions (MACS2 q < 0.05) for TCF7L2 ChIP-seq in Lgr5+ ISCs and the Crypt, and GATA4 in the intestinal epithelium.

**Supplementary table 2**. Genomic coordinates of peak-called regions (MACS3 q < 0.05) of accessible regions in single cell ATAC for each identified intestinal cell type.

**Supplementary table 3**. List of genes identified in Peak-to-gene-linkage by GREAT for TCF7L2 ISC & Crypt peaks and GATA4 peaks.

**Supplementary table 4**. qPCR primers & GEO accessions of published data used in this study. Note that the ChIP-seq data was realigned and reanalysed from raw fastq files and the single cell ATAC was reanalysed from published fragment files.

**Supplementary Figure 1.**
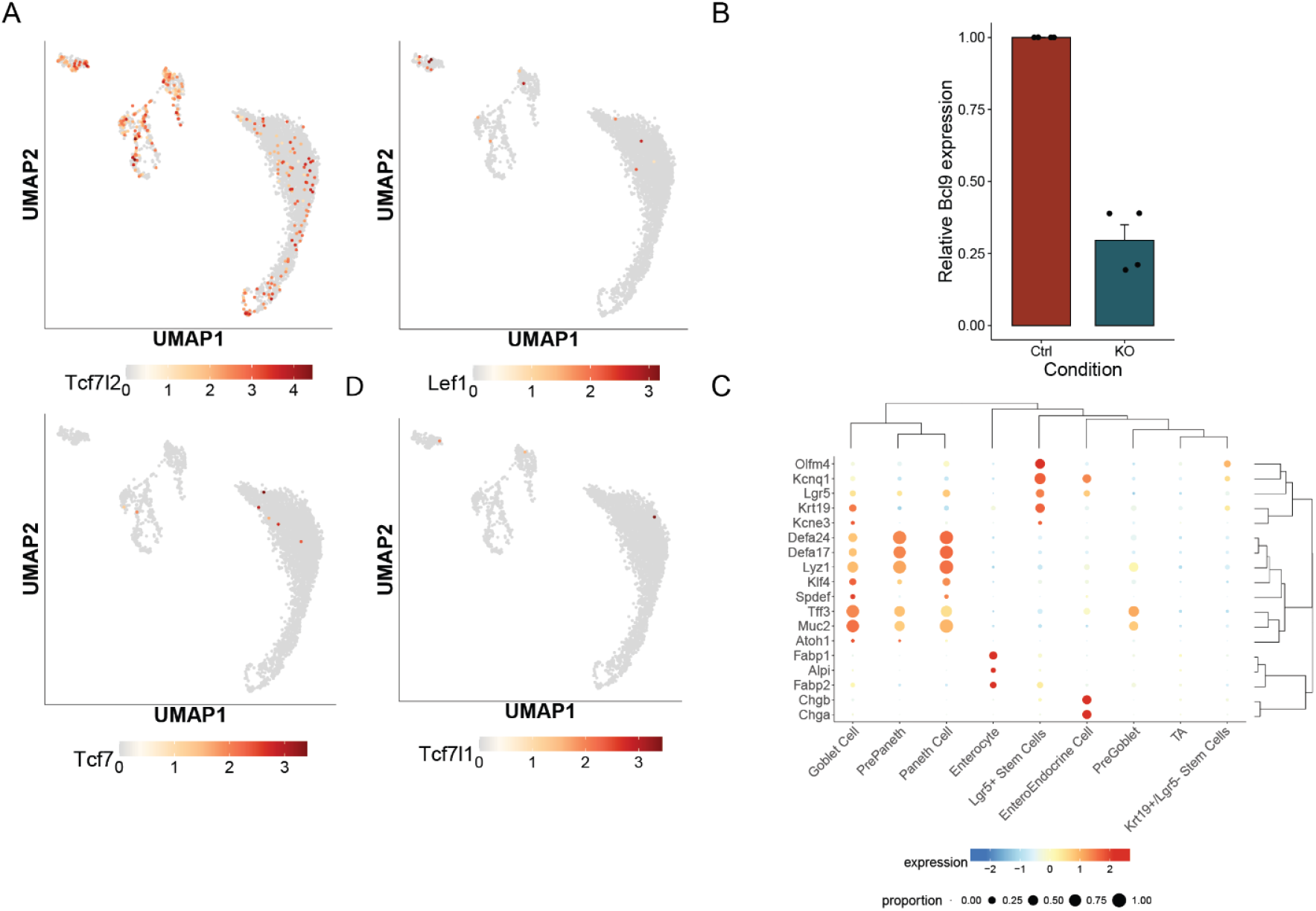
A) scRNA-seq expression of Tcf/Lef in intestinal organoids as overlay on an UMAP plot. B) RT-qPCR of Bcl9/9l 72h post-exposure to 1 μM 4OHT to determine knockout efficiency in intestinal organoids. C) Expression of marker genes used to annotate clusters displayed in a dotplot. Marker genes are based on PangloaDB^55^.

**Supplementary Figure 2.**
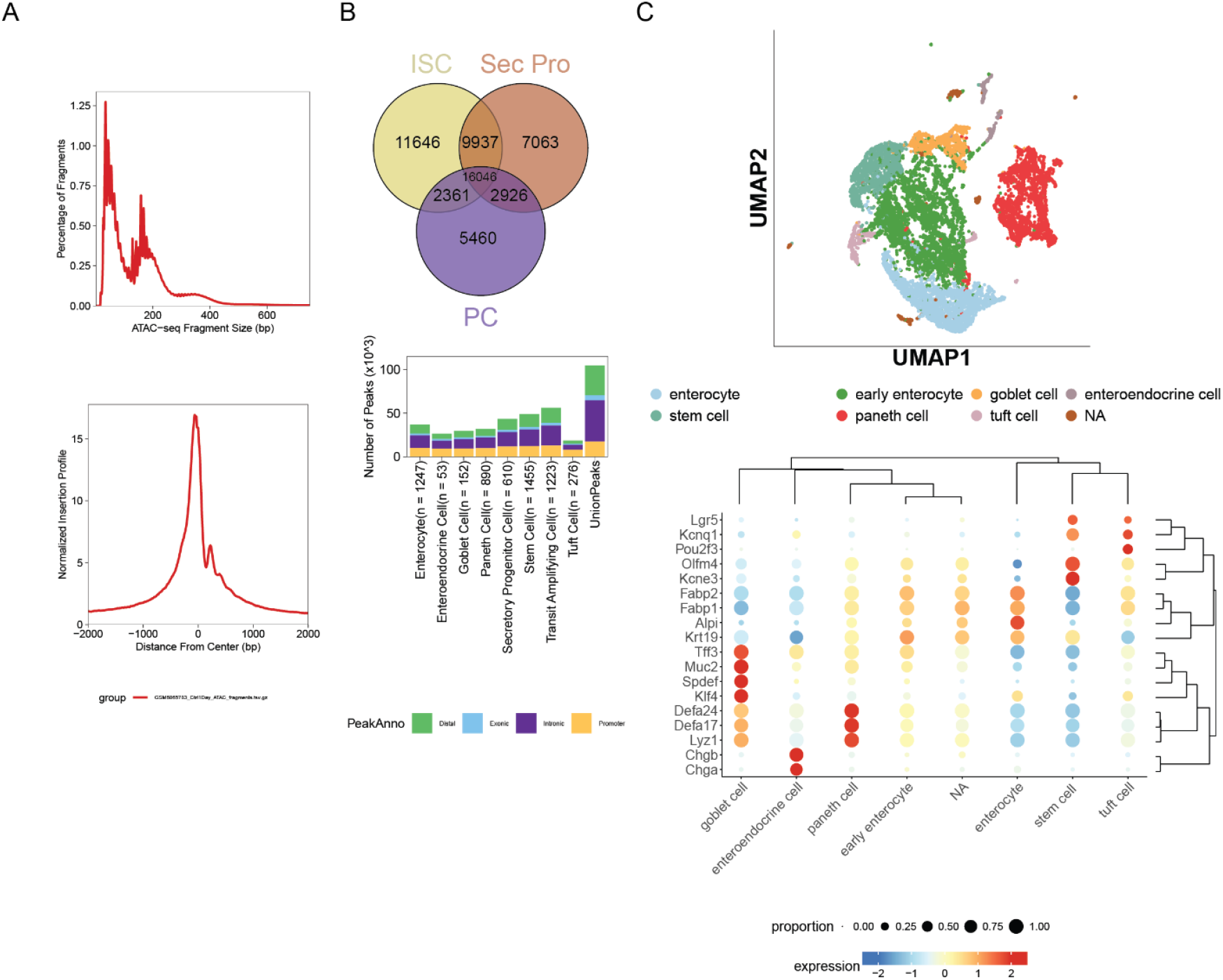
A) Quality control metrics scATAC-seq. B) Venn diagram displaying the overlap between pseudobulk ATAC peak called regions in Stem cells, Secretory progenitors and Paneth Cells (top). Genomic region annotation of pseudobulk ATAC peak called regions across all clusters (bottom). C) UMAP plot of scRNA-seq data used to annotate ATAC clusters (top). Expression of marker genes used to annotate clusters displayed in a dotplot. Marker genes are based on PangloaDB (bottom).

**Supplementary Figure 3.**
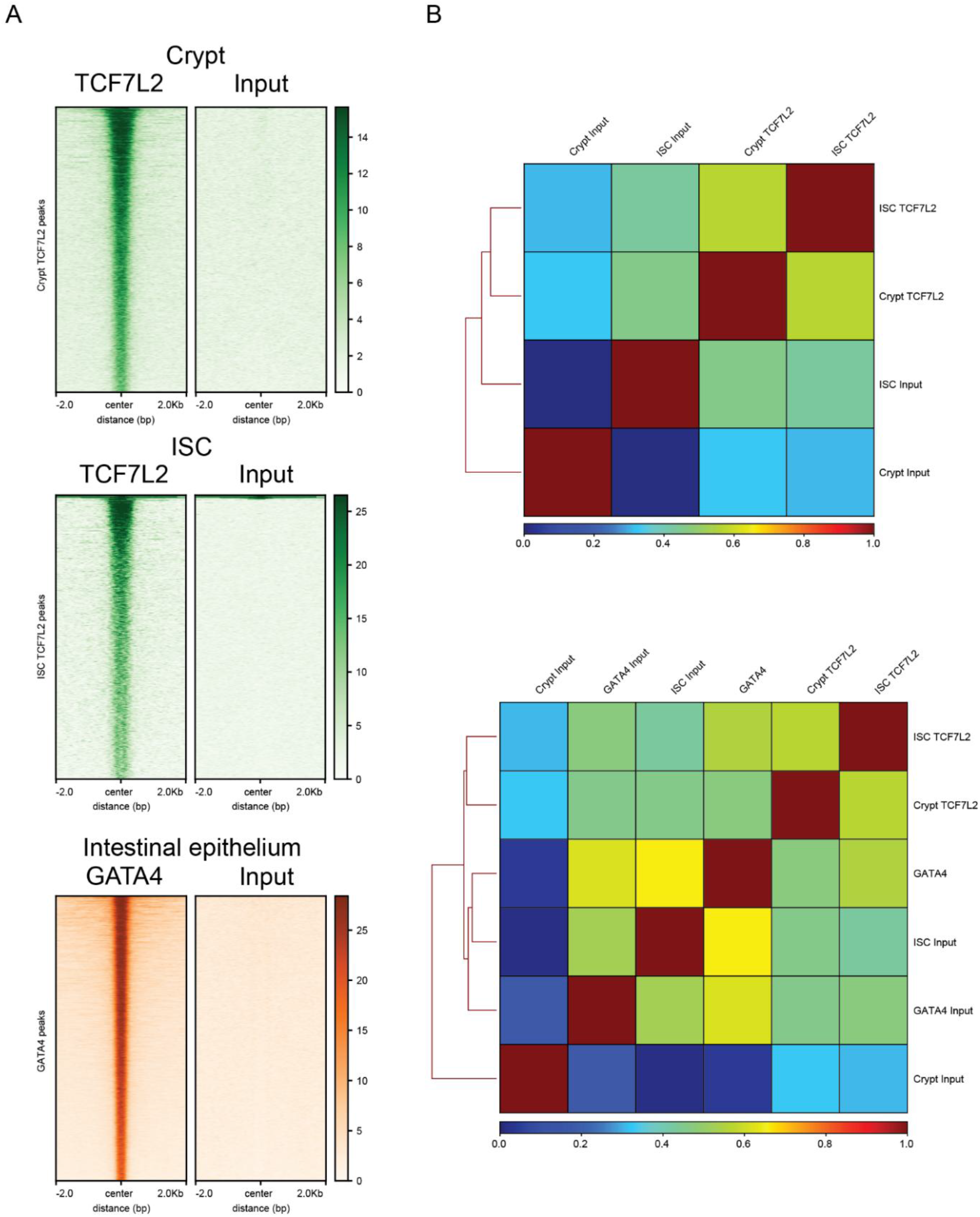
A) Signal intensity plots of both ChIP-seq signal and Input signal within peak called regions +/- 2kb for all ChIP-seq datasets. B) Overall data patterns shown by Spearman correlation.

